# Dual-stage inhibition of *Plasmodium falciparum* by a *Skeletocutis* derived fungal metabolite targeting Pyruvate Kinase II

**DOI:** 10.64898/2026.03.16.712130

**Authors:** Lucille Hervé, Nadia Amanzougaghene, Séverine Amand, Magali Blaud, Romain Coppée, Zaineb Fourati, Jean-François Franetich, Quentin Goor, Sandrine Houzé, Murielle Lohezic, Mathilde Patat, Véronique Sarrasin, Emilie Zelie, Valérie Soulard, Stéphane Mann, Anaïs Merckx

## Abstract

*Plasmodium falciparum* resistance to current first line treatments is threatening at-risk populations and underscores the urgent need for novel therapeutic targets and drugs. *P. falciparum* pyruvate kinases I and II are two essential enzymes with distinct roles and subcellular localizations within the parasite. *Pf*PyrKI is cytosolic, while *Pf*PyrKII is found in the apicoplast, a specific organelle of *Apicomplexa*, where it is required for the production of (d)NTPs essential for apicoplast maintenance. We identify skeletocutin E, a Basidiomycete-derived metabolite, as a specific inhibitor of *Pf*PyrKII. Skeletocutin E inhibits *in vitro* the activity of *Pf*PyrKII with an IC_50_ of 0.52 ± 0.08 µM through a mixed inhibition mechanism and does not affect the activities of three human pyruvate kinases. Structure–activity relationship analyses using synthetic skeletocutin E analogues allowed us to identify the molecular determinants of this inhibition. Furthermore, determination of the quaternary structure of *Pf*PyrKII by mass photometry, showed that this enzyme exists as monomers, dimers, and tetramers in equal proportions, revealing its singularity compared to other pyruvate kinases. Interestingly, skeletocutin E does not alter the distribution of the complexes, indicating that it does not interact at the subunit interfaces. Importantly, skeletocutin E inhibits *P. falciparum* growth in both blood and liver stages, with IC₅₀ values of 3.56 ± 0.50 µM in red blood cells and 3.70 ± 0.74 µM in primary human hepatocytes. Together, these findings establish *Pf*PyrKII as a druggable antimalarial target and identify skeletocutin E as a promising lead compound for the rational development of dual-stage antimalarial therapies.

## Introduction

Malaria is a major public health burden worldwide, with 610,000 deaths in 2024, mostly among children under five years old (1). This parasitosis is caused by *Plasmodium*, from the *Apicomplexa* phylum, with *P. falciparum* being the most lethal species. In its human host, *Plasmodium* undergoes two phases of intense multiplication, first in the liver, where the parasite develops within the hepatocytes, and then in erythrocytes, where parasite replication is associated with the symptoms of the disease. Resistance to the frontline treatments, which consists of artemisinin-based combination therapies, leading to treatment failures, has been reported since 2009 in the Greater Mekong Region (2). Since then, parasite mutations associated with artemisinin resistance have been observed in east African countries (3), threatening this continent where most malaria deaths occur. Therefore, identifying novel therapeutic targets is essential to expand the antimalarial drug pipeline with compounds whose mechanisms of action are unlikely to be affected by cross-resistance to existing treatments.

Pyruvate kinases (PyrK) catalyse the phosphorylation of ADP into ATP, converting phosphoenolpyruvate (PEP) into pyruvate, thus participating in glycolysis. In bacteria and lower eukaryotes, pyruvate kinase exists as a single form, although many bacteria and plants possess two isoenzymes. Vertebrates, by contrast, have four distinct isoenzymes (4). PyrK have proven to be therapeutic targets in number of diseases. Examples include the restoration of human PyrK deficiency in red blood cells, treated with Mitavipat (5–7), or the inhibition of a PyrK in the resistant bacteria *Staphylococcus aureus* (8) or in the parasite *Cryptosporidium parvum* in infected mice (9). Despite a few exceptions, PyrK enzymes are mainly active in a homotetrameric form (10–12). They are composed of four domains, with a catalytic pocket between the A and B domains, which is well-conserved across species, and an allosteric site that is distant by 40 Ångströms (Å) away from the catalytic site. PyrKs undergo transconformation between a tense (T) state and a relaxed (R) state, in which the catalytic pocket is closed. Their activity is regulated by various small molecules, including amino acids and metabolites. For example, phosphoenolpyruvate (PEP) acts as an allosteric activator, while fructose-1,6-bisphosphate, glucose-6-phosphate, and other sugars function as heterotropic activators (4, 13).

According to the literature, *P. falciparum* has two phylogenetically distant PyrKs: *P. falciparum* pyruvate kinase I (*Pf*PyrKI) which is cytosolic and glycolytic, and *P. falciparum* pyruvate kinase II (*Pf*PyrKII) which is compartmentalized and non-glycolytic (14–16). *Pf*PyrKI is essential for the parasite, particularly during the erythrocytic cycle when parasites are highly dependent on glucose (17). Its homotetrameric structure in both T- and R-states has been solved by X-ray crystallography (18, 19). *Pf*PyrKII is localized in the apicoplast, an essential organelle specific to *Apicomplexa* (16). *Pf*PyrKII is not nucleotide-specific and can phosphorylate any ribonucleotides or deoxyribonucleotides 5’ di-phosphates (d)NDP, providing ribonucleotides or deoxyribonucleotides 5’ tri-phosphates (d)NTP (15). Within the apicoplast, *Pf*PyrKII provides the necessary (d)NTP products required for DNA replication, transcription, translation, as well as for Fe-S clusters, fatty acids, and isoprenoid precursor pathways. Disruption of the *Pf*PyrKII gene or mislocalization of the protein disrupts the apicoplast, leading to parasite death (15). Because of its origin from two endosymbiotic events, the apicoplast has four membranes, and some pathways resembling those of plants or bacteria (20). Consequently, the apicoplast is therefore considered as a good therapeutic target, and can be targeted by antibiotics such as doxycycline, fosmidomycin, and azithromycin (21–23). Targeting the apicoplast can result in a delayed death phenotype, where the parasite can complete an erythrocyte cycle, but cannot divide at the end of the second. Delayed parasite death occurs when apicoplast maintenance is disrupted, as observed with doxycycline, which targets translation, whereas immediate death results from inhibition of the isoprenoid precursor pathway by fosmidomycin (24). However, despite the strong potential of the apicoplast as a therapeutic target, using antibiotics is not the most appropriate strategy in a world facing the emergence of multiresistant bacteria (21, 25). It is therefore necessary to identify other ways to interfere with the apicoplast biology and/or maintenance.

*Pf*PyrKII was classified as one of the 540 best therapeutic targets for which no inhibitors had been identified until recently (17, 26, 27). In this study, we sought to discover *Pf*PyrK inhibitors by screening molecules from a chemical library, two-thirds of which being compounds developed as part of malaria research. We tested the candidates *in vitro* using a pyruvate kinase enzymatic inhibition assay. This identified a specific *Pf*PyrKII inhibitor, which does not affect three human PyrK homologues, highlighting the specificity of *Pf*PyrKII as a therapeutic target. The mechanism of inhibition was characterized, and the molecular determinants of this interaction were investigated using analogues. Importantly, the compound inhibits *P. falciparum* growth in primary hepatocytes and in erythrocytes, without detectable toxicity to human cells. These results demonstrate a selective apicoplast-targeting *Pf*PyrKII inhibitor capable of blocking parasite growth at two life stages, supporting its potential for antimalarial drug development and possible applications against other pathogenic eukaryotes with a similar organelle.

## Results

### 1. *In vitro* kinetic propreties of recombinant *Pf*PyrK

Both *Pf*PyrKI and *Pf*PyrKII were purified as recombinant proteins from *Escherichia coli* expression system according to previous studies (15, 18), with final yields reaching 10-20 mg/L of bacterial culture. A six-histidine tag was fused to the N-terminus of *Pf*PyrKI for a first purification step by nickel affinity, followed by a size exclusion chromatography to obtain (His)_6_-*Pf*PyrKI (55 kDa). The first 41 residues of *Pf*PyrKII, containing the apicoplast targeting peptide were removed before cloning into the expression vector. An N-terminal MBP tag was used to purify MBP-*Pf*PyrKII with a first affinity step, and the MBP tag was then cleaved by the TEV protease prior to size exclusion chromatography leading to an untagged *Pf*PyrKII recombinant protein. We used an indirect enzyme assay, based on the formation of NAD^+^ from NADH, followed at 340nm, in the presence of lactate dehydrogenase, to confirm that both *Pf*PyrKI and *Pf*PyrKII were active *in vitro,* with catalytic constants (k_cat_) of 10.2 · 10^4^ ± 0.8 · 10^4^ min^-1^ for (His)_6_-*Pf*PyrKI and 1.1 · 10^4^ ± 0.3 · 10^4^ min^-1^ for untagged *Pf*PyrKII (Figure A1 & Table A1). These results enabled us to proceed with the enzymatic characterisation of both proteins.

(His)_6_-*Pf*PyrKI displayed sigmoid kinetics with respect to PEP and ADP with half-saturation constants (S_0.5_) of 0.289 ± 0.057 and 0.178 ± 0.028 mM, and Hill coefficients (n_H_) of 1.31 ± 0.15 and 1.22 ± 0.13, respectively, (Figure A1 A, A1 B & Table A1). These results indicated positive cooperativity and are consistent with the results previously published with the untagged *Pf*PyrKI (18). *Pf*PyrKII exhibited hyperbolic kinetics with respect to PEP, with an S_0.5_ of 0.059 ± 0.006 mM and an n_H_ of 1.13 ± 0.10 (Figure A1 C & Table A1). These results are consistent with the origin of *Pf*PyrKII, as plastid PyrK substrates are not allosteric activators. However, in ADP-related kinetics, the n_H_ was 1.29 ± 0.22 and S_0.5_ was 0.072 ± 0.011 mM, suggesting a possible allosteric activation by ADP and a specific affinity of *Pf*PyrKII for this substrate (Figure A1 D & Table A1). As shown by Swift et *al*., we found that *Pf*PyrKII could phosphorylate GDP with an S_0.5_ of 0.095 ± 0.014 mM and an n_H_ of 1.09 ± 0.15, indicating a lack of cooperativity for this substrate (Figure A1 E & Table A1). Our data showed a significantly higher affinity of *Pf*PyrKII for ADP than previously reported (0.072 ± 0.011 versus 0.25 ± 0.11 mM), leading to a better catalytic efficiency or k_cat_/S_0.5_ ratio (15,300mM^-1^min^-1^ versus 3,200 mM^-1^min^-1^). The differences were less pronounced for GDP, with a k_cat_/S_0.5_ ratio of 9,241 mM^-1^min^-1^ versus 5,890 mM^-1^min^-1^ in previous studies (15). Differences were also observed for the S_0.5_ relative to PEP (0.059 ± 0.006 versus 0.64 ± 0.05 mM), potentially due to variations in experimental procedures.

### 2. *Pf*PyrKII enzymatic activity is selectively inhibited by skeletocutin E

To have a control for inhibition of PyrK activity, suramin was selected and its ability to inhibit the enzymatic activities of *Pf*PyrKI and *Pf*PyrKII was studied. Suramin is an approved antiparasitic drug showing also applications against viruses and cancer that can inhibit *Tg*PyrKI, *Pf*PyrKI, and other PyrKs through competitive inhibition with ADP (18, 28). We confirmed in our hands that suramin inhibited *Pf*PyrKI, but with an IC_50_ of 7.02 ± 2.17 µM and a Ki of 3.22 µM. Additionally, suramin also inhibited *Pf*PyrKII, with an IC_50_ of 2.81 ± 0.772 µM and a K_i_ of 1.14 µM, indicating a stronger affinity for *Pf*PyrKII compared to *Pf*PyrKI (Figure A2). Suramin was used as a positive control and to validate inhibition assays.

In order to identify new putative *Pf*PyrK inhibiting molecules, we then tested compounds from a chemical library and a natural extract library of the French Muséum National d’Histoire Naturelle (MNHN), and involved them in an inhibition assay towards *Pf*PyrKI and *Pf*PyrKII enzymatic activities *in vitro*. Among the molecules from the library, the most effective inhibitor identified was MNHN-CH-0561, also known as skeletocutin E, synthetized at the MNHN in 1998 as an analogue of a Basidomycete metabolite (29). Skeletocutin E was then detected in Basidomycete *Skeletocutis* (30). It consists of a 14-carbon chain flanked at its both extremities by two citraconic anhydrides (Figure 1, compound **1**).

**Figure 1:**
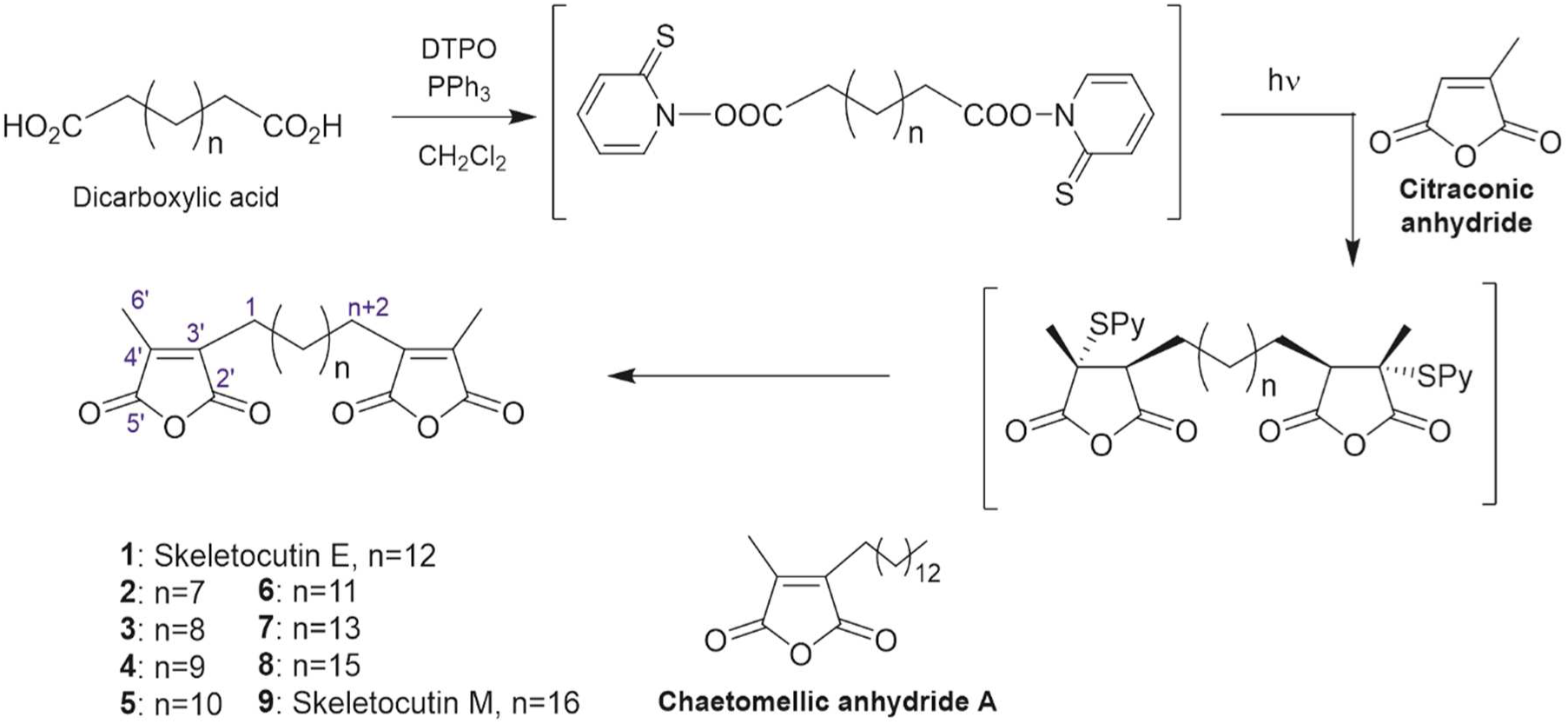
Synthesis of skeletocutin E and analogues as described by Poigny *et al.*(29) DTPO, 2,2’-dithiobis(pyridine N-oxide); Py, pyridyl. Atom numbering used for NMR assignment is indicated in blue.

In contrast, skeletocutin E showed only a weak inhibition of *Pf*PyrKI enzymatic activity, with a residual *Pf*PyrKI activity of 70.1 ± 7.92% at 30 µM, compared to residual *Pf*PyrKII activity of 10.9 ± 3.77% of at the same compound concentration (Figure 2A). Skeletocutin E inhibited *Pf*PyrKII whith ADP or GDP as substrates, with IC_50_ values of 0.52 ± 0.08 µM and 0.77 ± 0.08 µM, respectively (Figure 2B, D).

**Figure 2:**
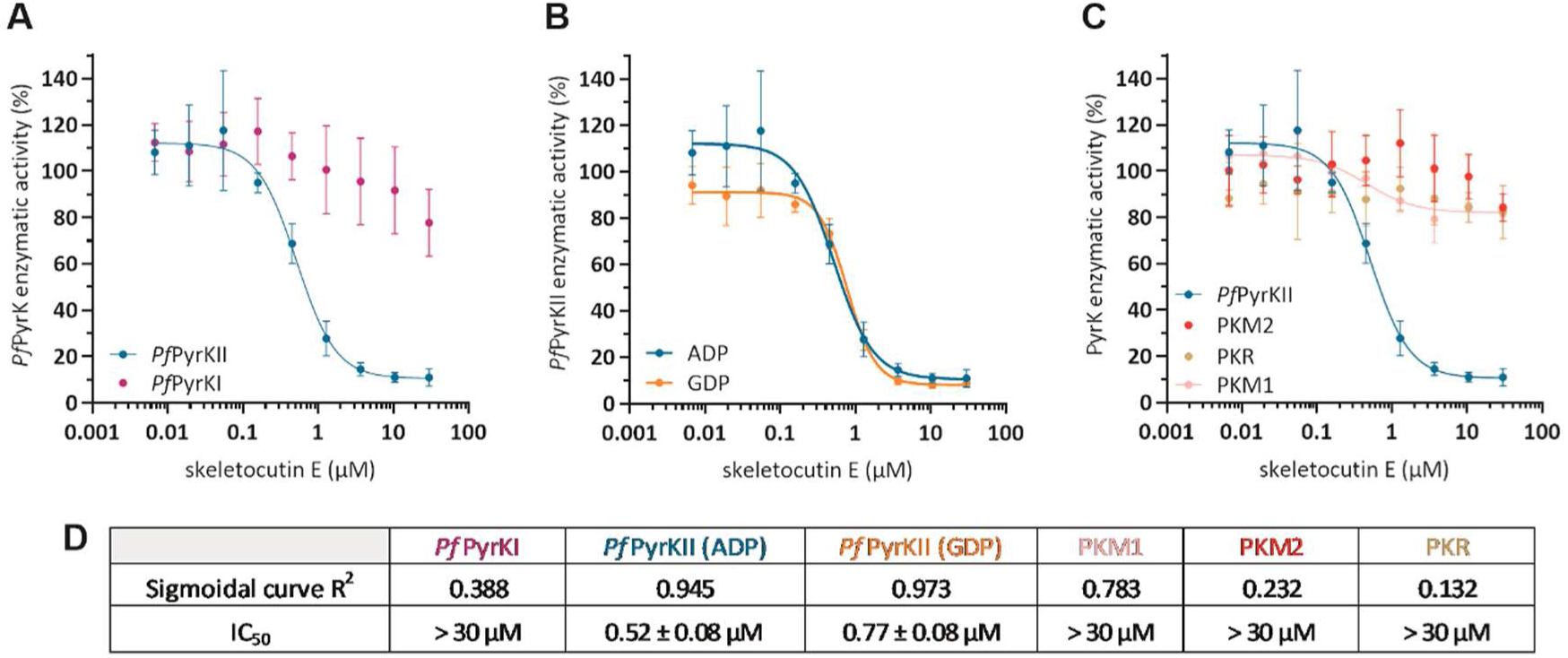
Skeletocutin E selectively inhibits *Pf*PyrKII. Enzymatic activity of *Pf*PyrKI (purple) and *Pf*PyrKII (green) normalized to the activity with 3% DMSO in the presence of a range of concentrations of skeletocutin E (values are means ± SD of 3 independent experiments) **(A)**. Enzymatic activity of *Pf*PyrKII normalized to the activity with 3% DMSO with substrate ADP (green) or GDP (orange) in the presence of a range of concentrations of skeletocutin E (values are means ± SD of 3 independent experiments) **(B)**. Enzymatic activity normalized to the activity with 3% DMSO in the presence of concentrations of skeletocutin E for *Pf*PyrKII (green), human PKM1 (pink), human PKM2 (red), and human PKR (beige) (values are means ± SD of 3 independent experiments) **(C)**. Coefficient of determination (R^2^) and IC_50_ related to each data set for a sigmoidal curve. A R^2^ close to 1 indicates a dose-dependent inhibition **(D)**.

Selectivity for the parasitic target is a prerequisite for any further development as an antimalarial drug candidate. To this end, inhibition assays were performed using three different types of commercially available human pyruvate kinases, PKM1 and PKM2 isoforms, and the red blood cells pyruvate kinase (PKR) (Table A1). Partial inhibition was observed for PKM1, as enzymatic activity followed a sigmoidal curve, with a remaining activity of 82.7 ± 2.74% in the presence of 30 µM skeletocutin E (Figure 2C, D). The enzymatic activities of PKR and PKM2 did not exhibit dose-dependent inhibition with skeletocutin E, showing 82.3 ± 11.5% and 84.2 ± 8.22% activity, respectively, at 30 µM skeletocutin E (Figure 2C, D).

Thus, skeletocutin E shows a high specificity for the *Pf*PyrKII enzyme and a high selectivity for its parasitic target, suggesting that this compound deserves further study to elucidate its mode of action.

### 3. Mechanism of *Pf*PyrKII inhibition by skeletocutin E

To analyse the mode of *Pf*PyrKII inhibition by skeletocutin E, we investigated the inhibition mechanism using Lineweaver-Burk plots. Skeletocutin E inhibited *Pf*PyrKII activity through a mixed inhibition mechanism (Figure 3A, B). The maximal initial velocity of *Pf*PyrKII is reduced by 4.30 nmol/min with 0.4 µM of skeletocutin E with respect to ADP. With the addition of 0.4 µM of inhibitor, the S_0.5_ of ADP decreased by 2.29-fold and the S_0.5_ of PEP decreased by 1.53-fold. Therefore, the influence of the molecule on the maximum velocity of the protein confirms that the inhibition of *Pf*PyrKII activity by skeletocutin E is not strictly competitive, and that the molecule interacts elsewhere than in the catalytic pocket. These results could explain the similar inhibition patterns when ADP or GDP are used as substrates. Also, these data are consistent with the fact that skeletocutin E is specific for *Pf*PyrKII since the amino acids of the PyrK catalytic site are well conserved among species, and thus one could expect that a specific *Pf*PyrKII inhibitor would not interact at this site. Finally, the mixed inhibition suggests that the inhibitor has different affinities for the enzyme alone and for the enzyme-substrate complex. For skeletocutin E, the affinity is lower for *Pf*PyrKII alone (K_ic_) than for the enzyme-substrate complex (K_iu_) (Figure A3). More specifically, this inhibitor binds to the PEP-*Pf*PyrKII complex with an affinity of 0.16 µM, and to *Pf*PyrKII without PEP with an affinity of 0.31 µM (Figure 3C). Similarly, it binds to the ADP-*Pf*PyrKII complex with an affinity of 0.11 µM, and to *Pf*PyrKII without ADP with an affinity of 0.42 µM (Figure 3D). Also, as revealed by the affinity constants, skeletocutin E has a higher affinity for *Pf*PyrKII complexed with substrates, which supports the existence of an interaction that is not in the catalytic pocket.

**Figure 3:**
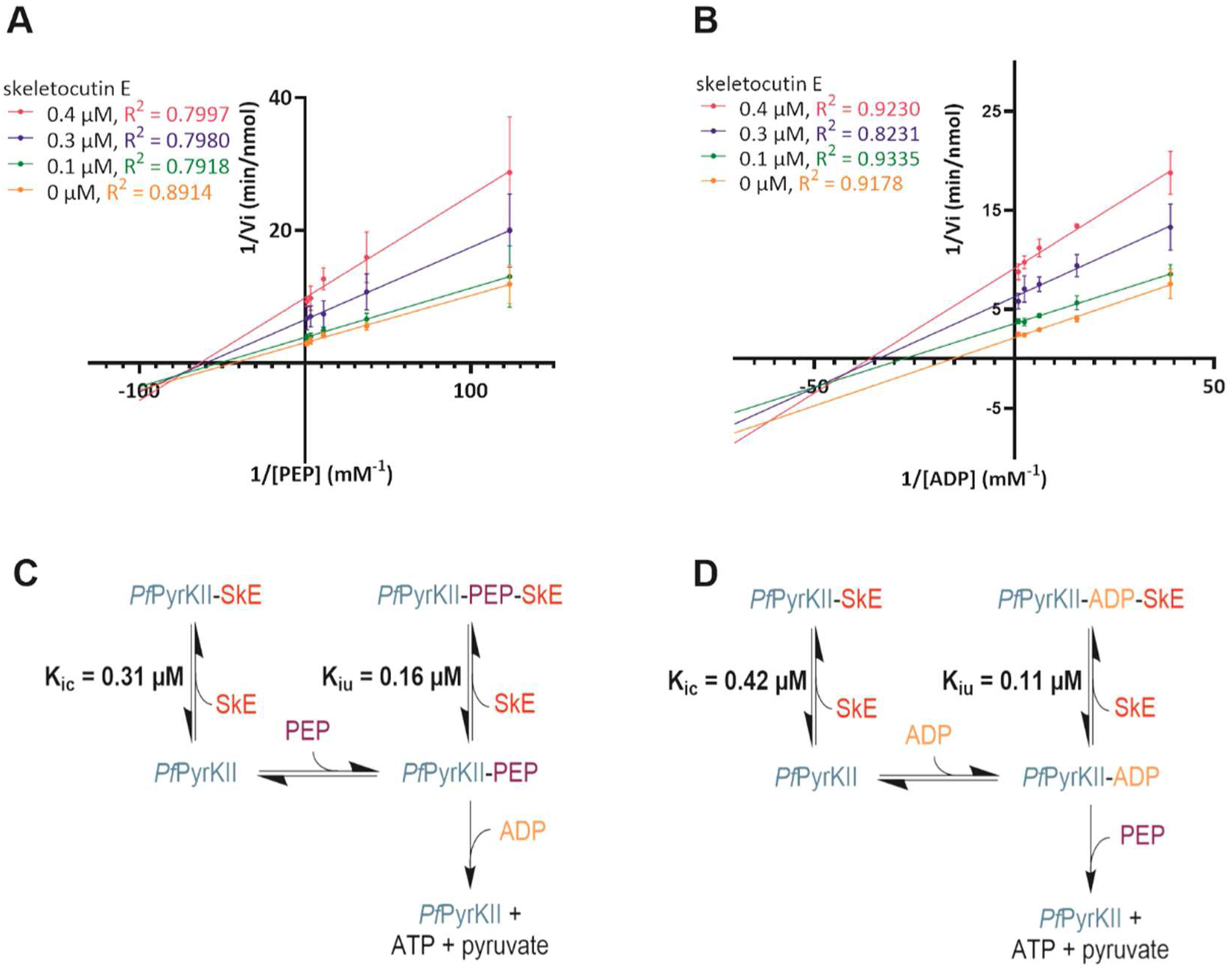
*Pf*PyrKII inhibitory mechanism. Lineweaver-Burk plots of *Pf*PyrKII for substrates PEP (**A**) and ADP (**B**) in the presence of 0, 0.1, 0.3, or 0.4 µM of skeletocutin E (values are means ± SD of 3 independent experiments). Schematic representation of the equilibrium between *Pf*PyrKII, its inhibitor skeletocutin E, and its substrate PEP (**C**) or ADP (**D**).

### 4. Quaternary organization of *Pf*PyrKII and its modulation by skeletocutin E

Since skeletocutin E does not bind the catalytic pocket, and no experimental 3D structure is available, we hypothesized that *Pf*PyrKII could form a homotetramer as most described pyruvate kinases. Therefore, we investigated the protein quaternary structure and observed the impact of the inhibitor interaction by mass photometry. The quaternary structures of *Pf*PyrKI and *Pf*PyrKII were analyzed in their respective enzymatic buffers. *Pf*PyrKI is homotetrameric (18, 19), and was used as a control. Mass photometry results show three *Pf*PyrKI oligomeric forms at 48, 158, and 399 kDa, with the latter being predominant and corresponding to the tetramer (Figure 4A). *Pf*PyrKI also appeared as a monomer (apparent molecular weight (aMW) = 48 kDa) and, to a lesser extent, as a dimer (aMW=158 kDa). Some inaccuracy in the calculated masses were observed due to differences in density between *Pf*PyrKs and the proteins used for calibration. Interestingly, *Pf*PyrKII exhibited a more dispersed distribution, with four major peaks at 63, 198, 328, and 446 kDa (Figure 4B). In comparison with the *Pf*PyrKI tetramer, we deduced that the peak at 446 kDa corresponds to the *Pf*PyrKII homotetramer. Therefore, the peaks at 63, 198, and 328 kDa correspond to the monomer, dimer, and trimer forms, respectively. Among the four quaternary structures, the monomer, dimer, and tetramer are equally represented, while the trimer is less abundant suggesting that the trimeric form might be a transition state between tetramer and dimer.

**Figure 4:**
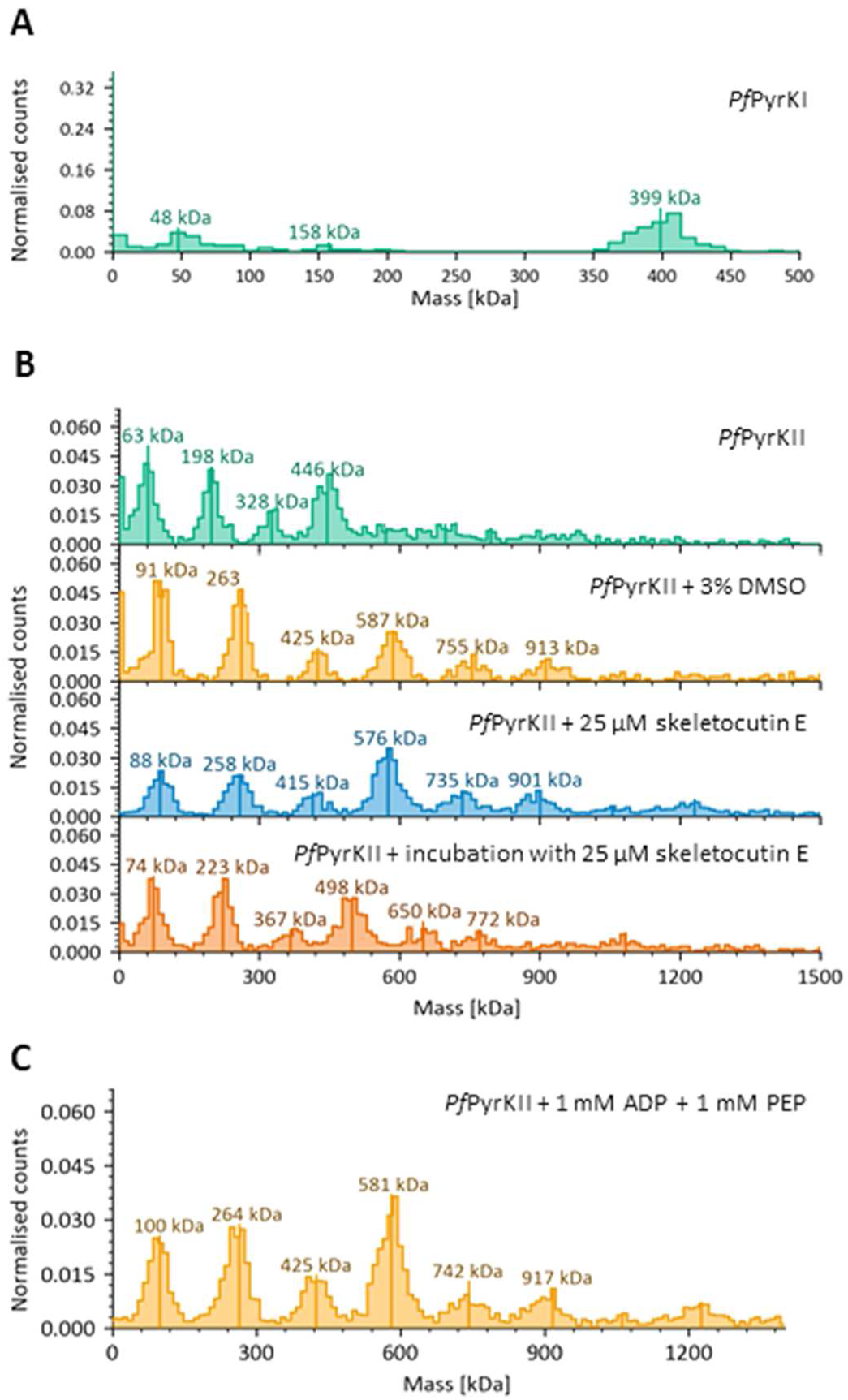
Analysis of the *Pf*PyrKII quaternary structure. Normalized count analyzed by mass photometry associated with *Pf*PyrKI **(A)**. Normalized counts analyzed by mass photometry associated with *Pf*PyrKII in the absence (*Pf*PyrKII) or presence of 3% DMSO and/or 25 µM of skeletocutin E with or without prior incubation between the protein and the molecule **(B)**. Normalized counts analyzed by mass photometry associated with *Pf*PyrKII in presence of 1 mM ADP and PEP **(C)**.

Upon addition of 3% DMSO, all *Pf*PyrKII forms shifted to higher masses of 91, 263, 425, and 587 kDa (Figure 4B). More defined peaks also appeared at 755 and 913 kDa, which could correspond to pentamers and hexamers. However, the distribution between monomer, dimer, trimer, and tetramer was not affected by DMSO. With the addition of 25 µM inhibitor, the distribution between the quaternary structures of *Pf*PyrKII remained dispersed, with monomer, dimer, trimer, tetramer, pentamer, and hexamer forms, with the proportion of monomer and dimer forms decreasing slightly in favour of the tetramer (Figure 4B). Incubation of the inhibitor with *Pf*PyrKII prior to analysis yielded distributions similar to those of the DMSO-free control, with a return to lower masses (74, 223, 367, and 498 kDa) (Figure 4B). These results suggest that DMSO is not responsible for the shift towards higher masses, and that neither DMSO nor the inhibitor have a significant impact on the distribution of the quaternary structure of *Pf*PyrKII. This implies that skeletocutin E does not disrupt amino acid interactions at the interface between subunits. To understand how the equilibrium between these four forms of *Pf*PyrKII is regulated, ADP and PEP substrates were added to the experiment. The peak masses shifted similarly to those observed with DMSO, shifting to 100, 264, 425, and 581 kDa, with no form being favored (Figure 4C). Therefore, substrate binding did not alter the equilibrium between the different forms of *Pf*PyrKII.

Despite the experimental error implying a difference in apparent masses, the mass photometry results show an unusual quaternary organization for *Pf*PyrKII, which is not disturbed significantly by the inhibitor or reaction substrates. This observation confirms that skeletocutin E does not interfere with the interfaces between subunits.

### 5. Inhibition of *Pf*PyrK2 by analogues

To define the molecular determinants of skeletocutin E (compound **1**) activity on *Pf*PyrKII, we designed a series of skeletocutin E analogues (Figure 1) and assessed their effects on *Pf*PyrK activity. The citraconic anhydride on its own did not show any inhibition of *Pf*PyrKII activity, indicating that the inhibition is not due to the cycle alone (Figures 1 & 5A). Furthermore, chaetomellic anhydride A (31), having a single citraconic anhydride linked to the aliphatic chain, lacked inhibitory activity, suggesting that two rings separated by a chain are essential for the observed effect (Figures 1 & 5A, E). To determine the optimal chain length for activity, several analogues of skeletocutin E with shorter (compounds **2-6**) and longer chains (compounds **7-9**) were synthesized (Tables A2 & A3) following the initial protocol for skeletocutin E (29), and their inhibitory effects were assessed *in vitro* on *Pf*PyrKI and *Pf*PyrKII activities. *Pf*PyrKI enzymatic activity was not significantly affected by the analogues, staying between 75 and 85 % (Figures 1 & 5B). According to our results, shortening the length of the carbon chain decreased the inhibitory potential of the molecule on *Pf*PyrKII (Figures 1 & 5B). The IC_50_ shifted to more than 10 µM with four carbons less than skeletocutin E in the chain. It even shifted to higher concentrations with one less carbon than skeletocutin E (Figure 5C, E). Conversely, longer carbon chains were not corelated to IC_50_ differences in the *Pf*PyrKII inhibition (Figure 5D, E). Together, these results indicate that the compound with optimal inhibition on *Pf*PyrKII is made of two cycles separated by a chain length of at least 14 carbons.

**Figure 5:**
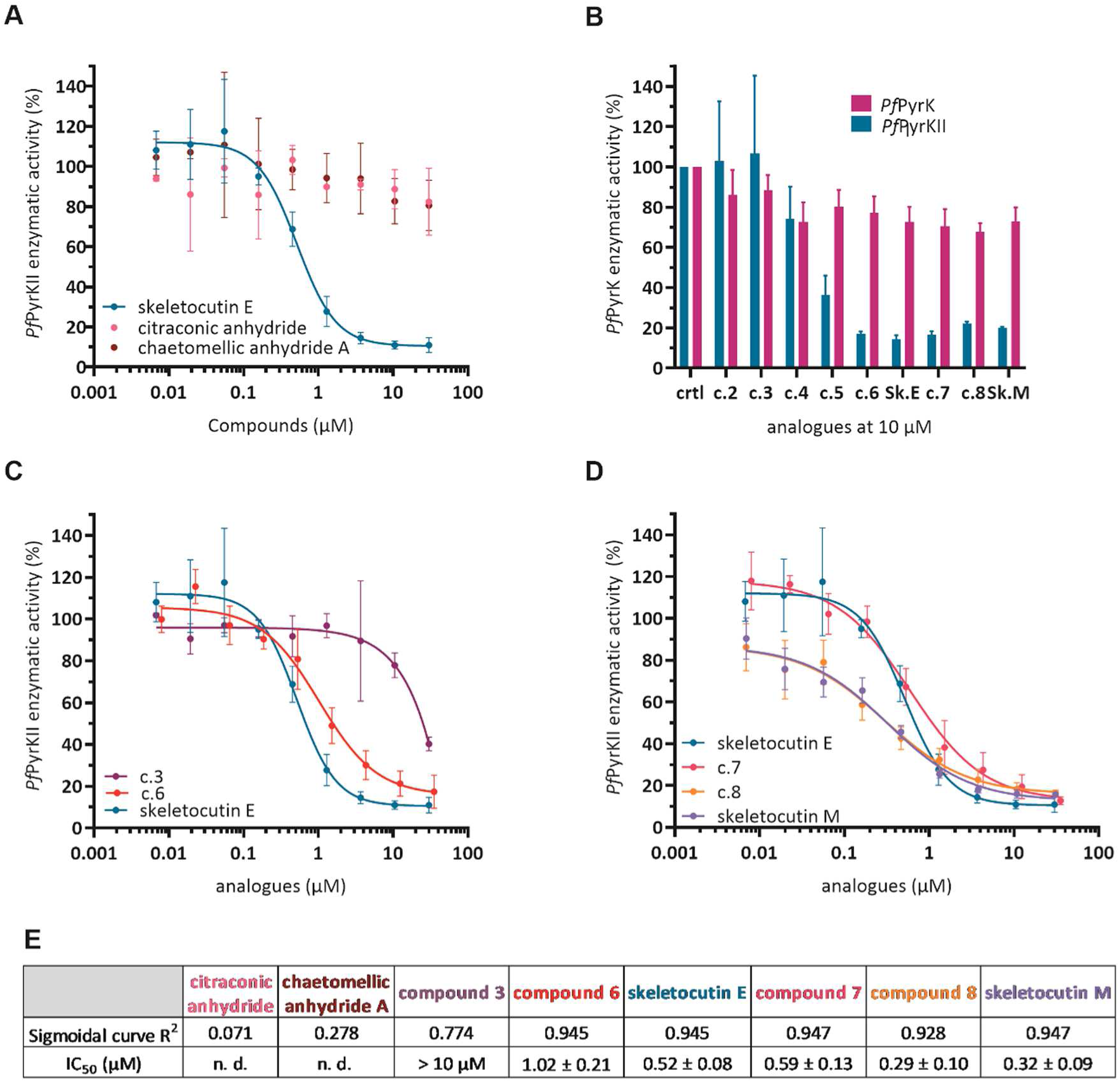
Inhibition of *Pf*PyrK enzymatic activity by skeletocutin E analogues *in vitro*. Enzymatic activity of *Pf*PyrKII normalized to activity with 3% DMSO in the presence of a range of concentrations of the citraconic and chetomellic A anhydrides (values are means ± SD of 3 independent experiments) (**A**). Enzymatic activity of both *Pf*PyrK normalized to the activity with 3% DMSO in the presence of 10 µM of the synthesized analogues (values are means ± SD of 3 independent experiments) (**B**). Normalized *Pf*PyrKII enzymatic activity of in the presence of a range of concentrations of synthesized analogues with a shorter (**C**) or longer (**D**) carbon chain (values are means ± SD of 3 independent experiments. Coefficient of determination (R^2^) and IC_50_ related to each data set for a sigmoidal curve. A R^2^ close to 1 indicates a dose-dependent inhibition (**E**).

### 6. Inhibition on *P. falciparum* growth in human hepatocytes

The activity of skeletocutin E on *P. falciparum* liver stages was assessed *in vitro* using primary human hepatocytes (PHH) infected with sporozoites. Cells were treated at the time of parasite inoculation to the PHH and for 6 days, with skeletocutin E concentrations ranging from 0.6 to 40 µM. As shown in Figure 6A, skeletocutin E inhibited liver-stage parasite development in a dose-dependent manner, leading to the almost complete elimination of the intracellular parasites (also called exoerythrocytic forms, EEFs) at a concentration of 40µM. Based on these results, the IC_50_ of skeletocutin E on *P. falciparum* liver stage *in vitro* was of 3.7 ± 0.74 µM (Figure 6C), with a decrease in parasite number of 59.15 ± 4.8% at 5µM of skeletocutin E compared to DMSO-treated and untreated controls (Figure 6A, C). This decrease in number was accompanied by a decrease in parasite size for the remaining EEFs (Figure 6B). Importantly, skeletocutin E was not toxic to PHH *in vitro* in the same range of skeletocutin E concentrations (Figure A4A). Four of the Skeletocutin E analogues tested on recombinant *Pf*PyrKII enzymatic activity were also evaluated for liver-stage inhibition: a single citraconic anhydride, chaetomellic anhydride A, and two chain-length variants (**3** and skeletocutin M). All were non-toxic to PHH (Figure A4B-E) and consistent with their weak or absent *Pf*PyrKII inhibition, the first three of them showed no effect on parasite development (Figure 6D-F). Surprisingly, skeletocutin M **9**, despite inhibiting *Pf*PyrKII similarly to skeletocutin E, did not impair parasite development within PHH (Figure 6G).

**Figure 6:**
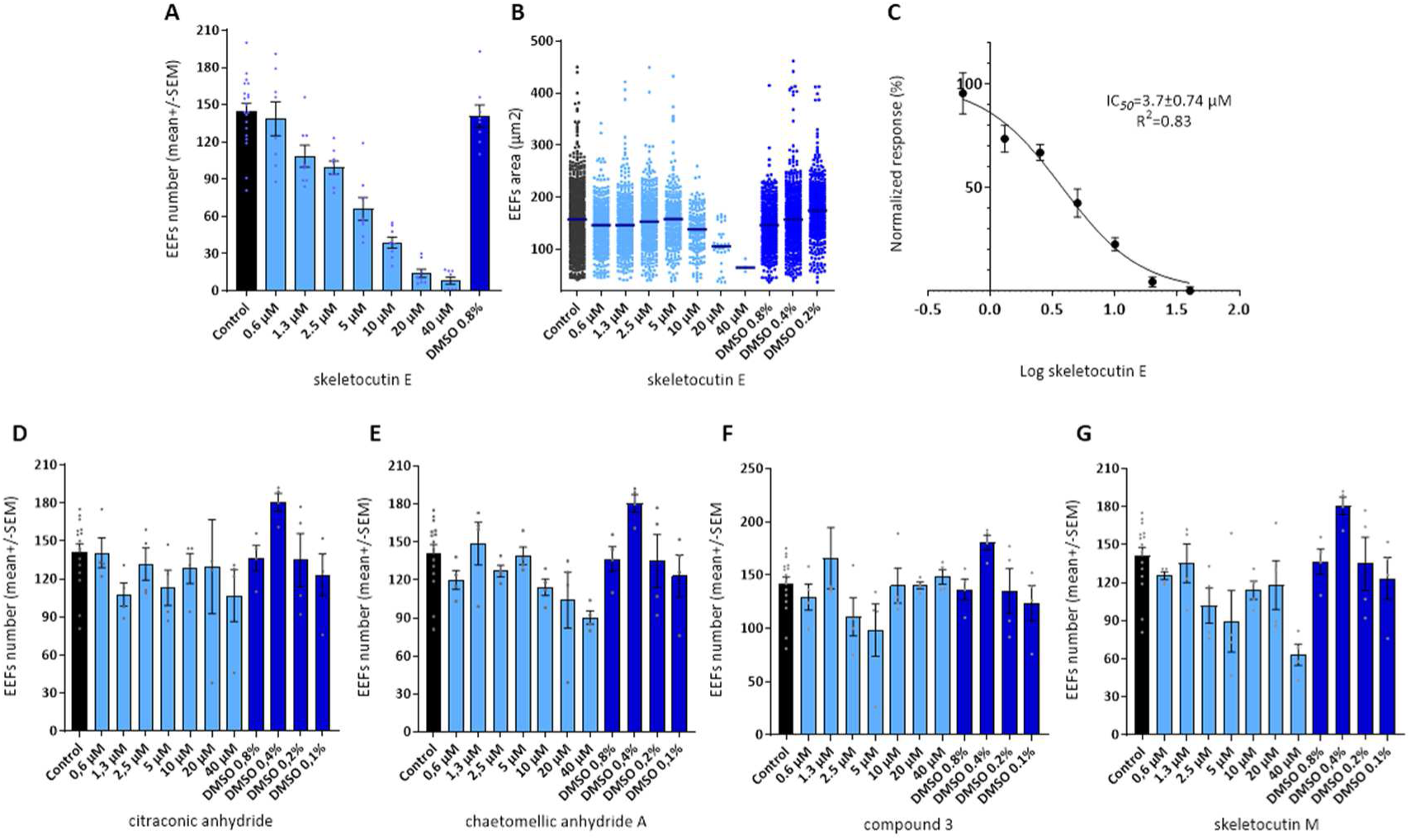
Effects of skeletocutin E and its analogues on *P. falciparum* development in hepatocytes. *Pf*HSP70 mouse antiserum was used to detect parasite schizonts. *P. falciparum* liver stage growth showed susceptibility to skeletocutin E as parasite numbers (**A**) and size (**B**) were reduced in a dose-dependent manner. The number of schizonts are expressed as mean ± SEM of four independent culture wells per condition. Parasite size values are derived from individual parasite measurements pooled from four independent culture wells per condition and expressed as surface area (μm²). Estimated IC₅₀ value for skeletocutin E against *P. falciparum*. Results from two experiments with four technical replicates each (**C**). Effect of citraconic anhydride (**D**), chaetomellic anhydride A (**E**), compound 3 (**F**) and skeletocutin M (**G**) on the number of *P. falciparum* schizonts, expressed as mean ± SEM of four independent culture wells per condition. In all experiments, solvent control (DMSO) correspond to the DMSO dilutions used for the respective compound dilutions (0.8%, 0.4%, 0.2%, 0.1%), matching the compound concentrations of 40, 20, 10, and 5 µM.

### 7. Inhibition of *P. falciparum* growth in human erythrocytes

Skeletocutin E was then tested for its activity on the development of *P. falciparum* within human erythrocytes, along with chloroquine as a positive control, in the conditions that are used for the culture of the erythrocytic stages. In these experimental settings, no inhibition of *P. falciparum* growth in red blood cells was observed (Figure 7A). Then, we hypothesized that serum components, particularly albumin, might interfere with skeletocutin E activity. To overcome this, infected cells were briefly exposed (6 h) to skeletocutin E (0.3–40 µM) in serum- and albumin-free medium, prior to culture in complete medium for one erythrocytic cycle. Parasitaemia was assessed 48 hours later. No hemolysis was observed in uninfected erythrocytes, confirming absence of toxicity of skeletocutin E to human red blood cells. Under serum-free exposure conditions, skeletocutin E yielded a dose dependent parasite inhibition with an IC₅₀ of 3.56 ± 0.50 µM (Figure 7B). Notably, pyknotic parasites were observed in infected erythrocytes exposed to ≥2.5 µM, consistent with severe developmental arrest (data not shown).

**Figure 7:**
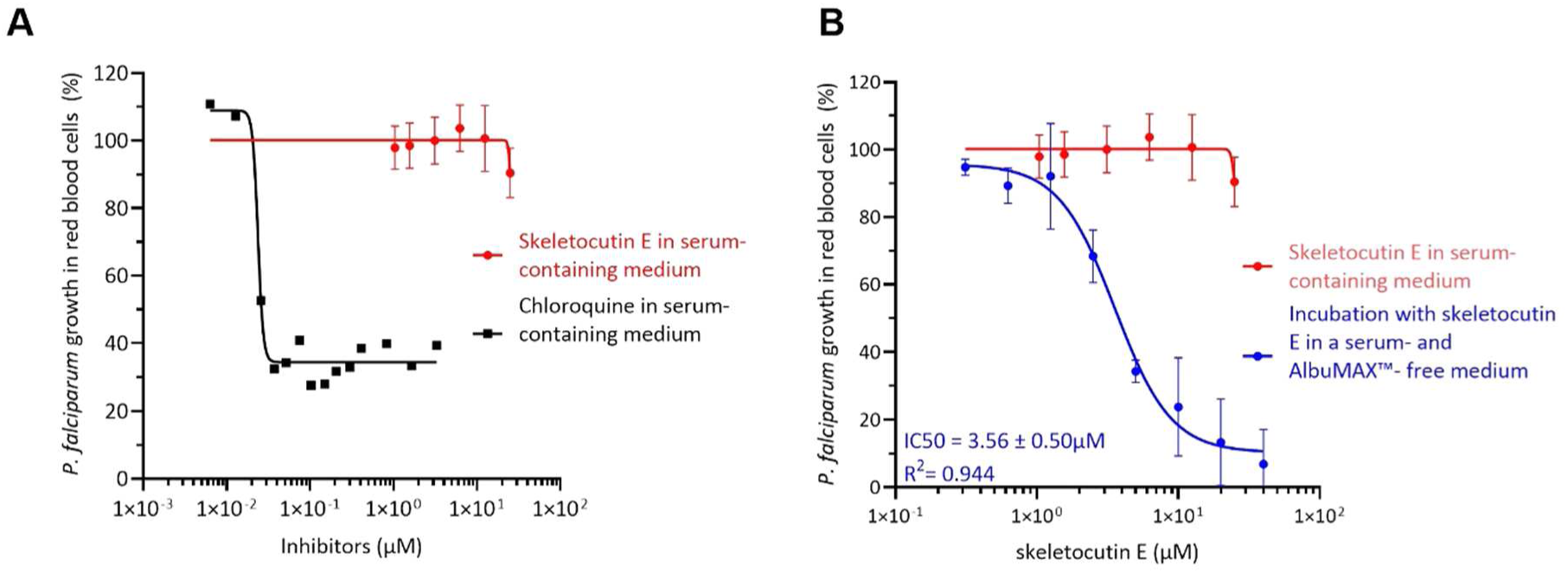
inhibition of of *P. falciparum* growth in infected red blood cells. Inhibition of *P. falciparum* growth in serum containing media with a range of chloroquine (black squares) as positive control of inhibition and skeletocutin E (red dots) (**A**). Inhibition of *P. falciparum* growth with skeletocutin E in serum containing media (red dots) and after a 6 h incubation in serum- and AlbuMAX™-free medium, before returning the parasites in serum and AlbuMAX™ containing medium (values are means ± SD of 3 independent experiments in duplicates) (**B**)

## Discussion

To combat malaria drug resistance, it is crucial to identify innovative targets, which, ideally, could be used to interfere with parasite development at multiple stages of its life cycle. In this context, targeting the apicoplast is a good strategy as it is essential in every stage of *Plasmodium*’s life cycle. During the liver stages, the parasite apicoplast is involved in heme and fatty acid synthesis (32–35), and during the erythrocytic stages, associated with malaria symptoms, the apicoplast supplies isoprenoid precursors (36, 37). Moreover, targeting the apicoplast can also lead to gametocyte death thereby blocking malaria transmission (36), and may potentially affect other pathogenic eukaryotes that harbor this organelle. Due to the endosymbiotic origin of the apicoplast, *Pf*PyrKII is phylogenetically far from mammalian PyrK and has singularities which may be exploited to be used as a therapeutic target. It is not specific with respect to nucleotide, and is larger than other PyrK (14, 15). Furthermore, we showed that *Pf*PyrKII exhibits a distinct quaternary structure distribution, with the homotetramers, dimers, and monomers present in roughly equal proportions, unlike other PyrKs.

In this study, we identified skeletocutin E as a potent and selective inhibitor of *Pf*PyrKII enzymatic activity which is far more effective than a previously reported inhibitor, procyanidin A1, which showed 90% inhibition of *Pf*PyrKII activity at 347 µM (27) compared to 10.5 µM for skeletocutin E. Moreover, skeletocutin E does not affect *Pf*PyrKI nor three of the human PyrK: PKM1, PKM2, and PKR. The only human PyrK that has not been tested in our study is PKL, a regulated enzyme localized in gluconeogenesis-producing tissues such as the liver, kidneys, and small intestine. However, PKL is an isoform of PKR, and skeletocutin E is not cytotoxic at 40 µM on hepatocytes (30), suggesting that it does not interfere with PKL activity.

Mechanistic characterization, including K_ic_ > K_iu_ values and mass photometry, suggests that skeletocutin E binds *Pf*PyrKII at a site distinct from the catalytic pocket or subunit interfaces. Although a mixed inhibition mechanism is supposed to imply potential binding of inhibitor both at the catalytic site and at an additional site of the enzyme, this remains controversial. Nevertheless, most experimental biases tend to yield K_iu_ > K_ic_; therefore, our observation of K_ic_ > K_iu_, which occurs in 10% of cases, indicates a lower likelihood of experimental error (38). Structural studies are thus needed to establish the precise interaction between *Pf*PyrKII and its inhibitor. No experimental structure of this enzyme is provided in the protein data bank or literature; it remains unsolved. The *Pf*PyrKII crystallogenesis was undertaken but unsuccessfully, therefore, we used an AlphaFold prediction instead (UniProtKB accession number Q8IJ37) (39). The prediction has a high confidence within the known pyruvate kinase structures: A, B, and C domains, whereas the N-terminal extension (residues 1 to 100), including the apicoplast targeting signal (14, 16), and two unstructured loops (residues 317 to 439 and 580 to 620), had very low confidence scores (plDDT < 50) (39). These unstructured loops are long insertions in compartmented PyrK enzymes of *Apicomplexa,* such as *P. falciparum*, *P. yoelii*, *P. berghei*, and *T. parva*, with unknown functions (14). Such unstructured loops are common in *Plasmodium* proteins and may contribute to crystallogenesis failure (40–42). The multimeric structure of *Pf*PyrKII was also predicted using AlphaFold3. In the tetrameric form, the second flexible loop observed in the monomer (residues 580 to 620) is much better structured in two α-helices in the center of the structure (Figure A5). These singularities of *Pf*PyrKII in comparison to other PyrK could be involved in the specificity of inhibition by the skeletocutin E.

Skeletocutin E is a secondary metabolite from a Kenyan wood-Inhabiting Basidiomycete: *Skeletocutis* (30). This molecule was extracted from *Skeletocutis* species, along with 17 analogues, including the reported metabolite tyromycin A (30, 43). Tyromycin A, found in another basidiomycete species: *Tyromyces lacteus*, differs from skeletocutin E by a two-carbon longer chain. It was identified as an inhibitor of leucine and cysteine aminopeptidases, and considered for immunomodulatory drug development. Thus, various synthesis pathways were described to produce these molecules and analogues, including skeletocutin E (29, 31, 44). However, subsequent studies did not reach the same conclusions regarding aminopeptidase inhibition (30, 43). These molecules and analogues were tested for biological activities on different organisms. Both skeletocutin E and tyromycin A inhibit *Bacillus subtilis* DSM10 strain with minimum inhibition concentrations of 44.8 µM and 21.0 µM respectively. Interestingly, both molecules were non-cytotoxic to mouse fibroblasts (under concentrations of 239 µM skeletocutin E and 224 µM tyromycin A) and HeLa (under concentrations of 88.4 µM skeletocutin E and 82.8 µM tyromycin A). These results are consistent with the non-inhibition of human PyrK by skeletocutin E, and confirm our finding that this molecule is not toxic to red blood cells and primary human hepatocytes. Additionally, skeletocutin E was not toxic to the human hepatocytes-derived carcinoma cell lines Huh-7.5 at 40 µM, and is antiviral against hepatitis C virus by reducing infectivity by more than 75% at this concentration (30). Taken together, these results indicate the potential of this molecule for its antimicrobial and antiviral activities, as well as its safety, as it is not toxic to four different types of mammalian cells.

In the present study, we observed an effect of skeletocutin E on the parasite at much lower concentrations than those reported for bacteria or the hepatitis C virus. This suggests a specific toxicity towards *P. falciparum*, possibly due to a particular interaction with *Pf*PyrKII, which differs from other PyrK enzymes. Interestingly, skeletocutin E inhibits two developmental stages of *P. falciparum*, a key feature for new antimalarial drug candidates. The IC₅₀ values of 3.7 ± 0.74 µM for the liver stage and 3.56 ± 0.50 µM for the red blood cell stage are of the same order of magnitude, which might indicate limited metabolization and similar accessibility to the target. For the liver stage, the requirement of *Pf*PyrKII activity has not yet been demonstrated, but the (d)NTPs it produces are probably necessary for DNA and RNA synthesis. Other pathways essential to liver stage-developing parasites may also depend on *Pf*PyrKII-derived ATP, such as translation, Fe–S cluster biogenesis, or isoprenoid precursor synthesis, as shown in red blood cells by Swift et *al*. (15). Furthermore, our results on the liver stage suggest that antiparasitic activity is highly dependent on the length of the compound’s carbon chain, with the optimal length being close to that of skeletocutin E. Future development of skeletocutin E should focus on enhancing bioavailability in human serum or albumin-rich environments, via rational modification or targeted delivery strategies. Thus, as a compound targeting a new therapeutic enzyme and inhibiting multiple *P. falciparum* developmental stages, skeletocutin E is an original and promising candidate that deserves further exploration for development as a novel antimalarial drug, potentially limiting the emergence of drug resistance.

## Materials and Methods

### Proteins expression and purification

*Pf*PyrK purification strategies were adapted from previously described protocols (15, 18). Genes of *Pf*PyrKI (PlasmoDB accession number PF3D7_0626800, residues 1-511) and *Pf*PyrKII deleted of its apicoplast signal (PlasmoDB accession number PF3D7_1037100, residues 42-745) were codon optimized and cloned in a pRSF-Duet and pMAL-c6T plasmids respectively. (His)_6_-*Pf*PyrKI and MBP-*Pf*PyrKII were expressed in *Escherichia coli* BL21-Codon Plus cells in Lysogeny Broth (LB) medium complemented with the appropriate antibiotic according to the plasmids. Protein expression was induced with 0.3 mM of isopropyl β-d-1-thiogalactopyranoside (IPTG) at 37 °C during 4 h for (His)_6_-*Pf*PyrKI; and at 30°C during 6h for MBP-*Pf*PyrKII. Cells were harvested by centrifugation then resuspended in lysis buffer (50 mM triethanolamine-HCl pH 7.2, 300 mM KCl, 20 mM imidazole, 10% glycerol for (His)_6_-*Pf*PyrKI; 20 mM Tris HCl pH 7.5, 200 mM NaCl with 3 µg/mL DNase for MBP-*Pf*PyrKII), complemented with 1 mg/mL lysozyme (Sigma) and cOmplete EDTA-free protease inhibitor cocktail (Roche), and lysed by sonication. The soluble fraction was collected by centrifugation (48 000 g, 30 min, 4°C) and purified by affinity chromatography. (His)_6_-*Pf*PyrKI was eluted with an imidazole gradient up to 500 mM and purified on a size exclusion chromatography Superdex S200 16/600 column (Cytiva) in a 20 mM triethanolamine-HCl pH 7.2, 50 mM KCl, 10 mM MgCl2, 10% glycerol buffer. MBP-*Pf*PyrKII was eluted with 100 mM maltose, before TEV protease cleavage (with 1:100 ratio and 1 mM 1,4-dithiothreitol), and size exclusion chromatography in a 20 mM Tris HCl pH 7.5, 200 mM NaCl buffer. Fractions containing the target proteins were concentrated and conserved at -80°C. Purified *Pf*PyrKI and *Pf*PyrKII were confirmed by mass spectrometry.

### *In vitro* pyruvate kinase enzymatic assays

*Pf*PyrK activity was measured using a coupled enzyme assay with lactate dehydrogenase (LDH) according to established protocols and buffers (15, 18). Oxidation of NADH was assessed at 340 nm with a SAFAS UVmc2® spectrometer every 30 sec for at least 10 minutes in 50 µL quartz cuvettes (Hellma Analytics). The exact concentrations of NADH, ADP and GDP (Sigma) were measured by their absorbance at 340, 259 and 253 nm, respectively. The PEP (Roche) solution was adjusted to pH 7, and its concentration measured with a PEP assay kit (Cell Biolabs). The *Pf*PyrKs were thawed and centrifuged (21,000 g, 10 min, 4°C) before being analysed by Bradford assay to measure *Pf*PyrK concentration, and diluted before each assay. *Human* PK were purchased from Sigma-Aldrich (Pyruvate Kinase M2 human cat# SAE0021; PKM1 active human cat# SRP0415; PKLR Var2 active human, cat#SRP0414). The reaction mixture contains buffer suitable for each PyrK from literature (15, 18, 45, 46) or the supplier’s recommendations, an appropriate PyrK concentration for the absorbance to be linear for few minutes, 0.2 mM NADH and 1.9 U of LDH (Roche). For enzyme kinetics analysis, the reaction mixture contains one of the substrates in excess (> 3**·** S_0.5_), and the reaction was initiated with a range of concentrations of the second substrate. For the enzymatic activity inhibition study, the reaction mixture contained PEP and the reaction was triggered with ADP or GDP, with all substrate concentrations equal to 1.5**·** S_0.5_. The composition of the buffer, the concentrations of PyrK and substrates are presented in the supplementary information (Table S4 & S5). All reactions were measured in a final volume of 70 µL, in duplicate or triplicate. For skeletocutin E and its analogues in DMSO, DMSO percentage is set to 3 %. The initial enzyme velocity was calculated from a tangent fitted to the reaction curve and, in inhibition assays, it was normalised to the initial velocity of the enzyme with 3 % DMSO. The secondary plots were fitted as sigmoids on GraphPad Prism and the kinetic parameters S_0.5_, V_max_, n_H_ and IC_50_ were extracted. For the inhibition mechanism, the kinetics parameters S_0.5_’ and V_max_’ were determined from the Lineweaver-Burk plot. The K_i_ values for competitive inhibition were calculated from IC_50_. For mixed inhibition, the K_i_ values were extracted from tertiary plots representing slope and y-intercept of Lineweaver-Burk as a function of inhibitor concentration. All secondary and tertiary plots are available in supporting information.

### Mass photometry

A mass photometry analysis was performed on a Refeyn TwoMP in a volume of 20 µL. All samples, substrates, and buffer (50 mM triethanolamine-HCl pH 7.2, 100 mM KCl, 10mM MgCl_2_ for *Pf*PyrKI, and 26.5 mM Tris-HCl pH 7.5, 50 mM KCl, 10 mM MgCl_2_ for *Pf*PyrKII) were centrifuged (21,000 g, 10 min, 4°C) to remove aggregates. Eighteen µL of buffer were placed on the laser, then 40 nM of *Pf*PyrKI or 50 nM of *Pf*PyrKII were added, mixed up and down with a micropipette, and counts were measured for a few minutes. For analysing the interaction with DMSO or skeletocutin E, both of them were include in the 18 µL of buffer and the percentage of DMSO were set to 3%. For incubation of *Pf*PyrKII and skeletocutin E, they were mixed together for few minutes at ambient temperature and added together to the buffer. The counts were analysed using Refeyn software and normalised to the total number of counts. The mass was estimated using BSA and urease calibration curves.

### Skeletocutin E and analogues synthesis

Skeletocutin E and analogues synthesis were achieved from dicarboxylic acids by double Barton radical decarboxylation, followed by condensation with citraconic anhydride in dry dichloromethane, as previously described (29) (Figure 1). Pure compounds **2-4** were isolated as colourless oils with yields of 42%, 31%, and 31%, respectively, while **5-9** and **1** were obtained as white powders with respective yields of 39%, 30%, 30%, 27%, 28%, and 36%. NMR and mass spectrometry characterizations of the molecules are reported in Tables A2 and A3.

### *P. falciparum* inhibition assays in erythrocytes

#### Inhibition assay in serum-containing medium

The *P. falciparum* 3D7 strain was cultured as previously described (47), in RPMI 1640 (with 5 mg/L phenol red), supplemented with 2.5 mM HEPES (pH 7.2-7.5), 2 mM L-glutamine, 0.05 mg/mL gentamicin, 10 % v/v human serum (PPA), 9 mM dextrose, 0.2 mM hypoxanthine. Parasites were synchronized at the ring stage using sorbitol. After treatment with sorbitol, cultures consisted mainly of ring-form parasites. A suspension was prepared with 2.5 % haematocrit and 0.3 % parasitaemia, to which 20 % v/v of tritiated hypoxanthine is added (>97 %, 5 mCi). Skeletocutin E solution was prepared by diluting the stock (2.5 mM in 50% DMSO) in water with 3% DMSO. In 96-well plates with filters (96 GF/C Microplates), 95 µL of parasite suspension was added to 5 µL of skeletocutin E or chloroquine solution. The final DMSO concentration was a maximum of 0.8%. Each condition was tested in triplicate. Plates were incubated for 42, 66, or 90 h at 37°C. After incubation, plates were frozen at -20°C, thawed, and dried for 2h. Scintillation oil was added, and radioactivity was measured for 2 min per well in a scintillation counter (MicroBeta2, Revvity), expressed in counts per minute

#### Inhibition assay with incubation in serum free medium

Infected erythrocytes were cultured in RPAS (RPMI-AlbuMAX™-Serum) medium (RPMI 1640 medium (Gibco), supplemented with 25 mM HEPES (Gibco), 2 mM L-glutamine (Gibco), 0.05 mg/ml gentamicin (Gibco), 2 % human serum and 0.5% of AlbuMAX™). Infected red blood cells were washed in culture medium without serum nor AlbuMAX™, and a suspension was prepared with 2.5% haematocrit and 1-2 % parasitaemia. A series of dilutions of skeletocutin E was prepared in DMSO, ranging from 39 µM to 5 mM. Five hundred µL of parasite suspension, in a medium without serum nor AlbuMAX™, was added to a volume of 4 µL of skeletocutin E or DMSO (control) solution in a 24-wells culture plate (TPP). The final DMSO concentration was 0.8%. Each condition was tested in duplicate, in three independent experiments. Plates were incubated in a gas mixture of 5 % O_2_, 5 % CO_2_ and 90% N_2_ at 37°C. After 6 h, skeletocutin E solutions were removed and 500 µL of fresh medium with 2% serum and 0.5 % AlbuMAX™, without compound, were added. Plates were replaced at 37°C in the same gas mixture for one parasite cycle. After incubation, smears were performed for all experimental conditions to assess parasitaemia. IC_50_ was derived from non-linear regression of a four-parameter logistic / Hill model.

### *P. falciparum* inhibition assays in hepatocytes

*P. falciparum* liver stage culture was performed as described in Ashraf et al (48). Briefly, NF54 *P. falciparum* sporozoites were obtained from the Centre d’Elevage et d’Infection des Anophèles (CEPIA, Institut Pasteur, Paris) and 30,000 sporozoites were inoculated to primary human hepatocytes (PHH) in 384-well plates. After a centrifuge at 560 g for 10 min at room temperature to allow for rapid sporozoite sedimentation onto hepatocytes, the cells were incubated at 37°C under 5% CO_2_. Three hours later, cultures were washed and a layer of matrigel (Corning Life Sciences) was added, as per the manufacturer’s recommendations. After 45 min at 37°C, culture medium was added and was thereafter renewed every 24 h. For compound testing, PHH cultures were exposed to a two-fold dilution drug series, from 40 µM to 0.6 µM, along with no-drug and solvent (DMSO) controls. The compounds tested were added at the same time as the sporozoites. Each experimental condition included four culture wells. Culture medium with or without drugs or DMSO was refreshed every 24 hours, and the development of liver stage parasites was stopped 6 days after infection, by fixation with paraformaldehyde (PFA) 4% for 15 minutes at room temperature. The parasites were stained using a anti-*Pf*HSP70 mouse immune serum, revealed with an Alexa 488-conjugated goat anti-mouse immunoglobulin (Molecular Probes). Host cell and parasite nuclei were labelled with 4′,6-diamidino-2-phenylindole (DAPI). EEFs numbers and sizes were measured using a Cell Insight High-Content Screening platform with Studio HCS software (Thermo Fisher) on the Celis Platform (ICM, La Pitié-Salpêtrière, Paris), as described previously (48). IC_50_ was derived from non-linear regression of a four-parameter logistic / Hill model.

## Acknowledgments

The authors acknowledge K. Callabro, A. Gand, S. Gillet, C. Maulay-Bailly, H. Munier-Lehmann & S. Rosario for scientific discussions, mass photometry experiments and the French National Chemical Library (http://chembiofrance.org) for providing the products for screening. This work has received support under the program “Investissement d’Avenir” launched by the French Government and implemented by ANR, with the reference « ANR-18-IdEx-0001 » as part of its program « Emergence ».

## References

1. World malaria report 2025. https://www.who.int/teams/global-malaria-programme/reports/world-malaria-report-2025. Retrieved 29 January 2026.

2. Dondorp AM, Nosten F, Yi P, Das D, Phyo AP, Tarning J, Lwin KM, Ariey F, Hanpithakpong W, Lee SJ, Ringwald P, Silamut K, Imwong M, Chotivanich K, Lim P, Herdman T, An SS, Yeung S, Singhasivanon P, Day NPJ, Lindegardh N, Socheat D, White NJ. 2009. Artemisinin resistance in Plasmodium falciparum malaria. N Engl J Med 361:455–467.

3. Balikagala B, Fukuda N, Ikeda M, Katuro OT, Tachibana S-I, Yamauchi M, Opio W, Emoto S, Anywar DA, Kimura E, Palacpac NMQ, Odongo-Aginya EI, Ogwang M, Horii T, Mita T. 2021. Evidence of Artemisinin-Resistant Malaria in Africa. N Engl J Med 385:1163–1171.

4. Enriqueta Muñoz Ma, Ponce E. 2003. Pyruvate kinase: current status of regulatory and functional properties. Comp Biochem Physiol B Biochem Mol Biol 135:197–218.

5. Rab MAE, Van Oirschot BA, Kosinski PA, Hixon J, Johnson K, Chubukov V, Dang L, Pasterkamp G, Van Straaten S, Van Solinge WW, Van Beers EJ, Kung C, Van Wijk R. 2021. AG-348 (Mitapivat), an allosteric activator of red blood cell pyruvate kinase, increases enzymatic activity, protein stability, and ATP levels over a broad range of PKLR genotypes. Haematologica 106:238–249.

6. Al-Samkari H, van Beers EJ. 2021. Mitapivat, a novel pyruvate kinase activator, for the treatment of hereditary hemolytic anemias. Ther Adv Hematol 12:20406207211066070.

7. Luke N, Hillier K, Al-Samkari H, Grace RF. 2023. Updates and advances in pyruvate kinase deficiency. Trends Mol Med 29:406–418.

8. Jia J, Luo Y, Zhong X, He L. 2022. Methicillin-Resistant Staphylococcus Aureus (MRSA) Pyruvate Kinase (PK) Inhibitors and their Antimicrobial Activities. Curr Med Chem 29:908–923.

9. Khan SM, Zhang X, Witola WH. 2021. Cryptosporidium parvum Pyruvate Kinase Inhibitors With in vivo Anti-cryptosporidial Efficacy. Front Microbiol 12:800293.

10. Pawluk A, Scopes RK, Griffiths-Smith K. 1986. Isolation and properties of the glycolytic enzymes from Zymomonas mobilis. The five enzymes from glyceraldehyde-3-phosphate dehydrogenase through to pyruvate kinase. Biochem J 238:275–281.

11. Plaxton WC, Smith CR, Knowles VL. 2002. Molecular and regulatory properties of leucoplast pyruvate kinase from Brassica napus (rapeseed) suspension cells. Arch Biochem Biophys 400:54–62.

12. Nairn J, Duncan D, Gray LM, Urquhart G, Binnie M, Byron O, Fothergill-Gilmore LA, Price NC. 1998. Purification and characterization of pyruvate kinase from Schizosaccharomyces pombe: evidence for an unusual quaternary structure. Protein Expr Purif 14:247–253.

13. Schormann N, Hayden KL, Lee P, Banerjee S, Chattopadhyay D. 2019. An overview of structure, function, and regulation of pyruvate kinases. Protein Sci Publ Protein Soc 28:1771–1784.

14. Chan M, Tan DSH, Sim TS. 2007. Plasmodium falciparum pyruvate kinase as a novel target for antimalarial drug-screening. Travel Med Infect Dis 5:125–131.

15. Swift RP, Rajaram K, Keutcha C, Liu HB, Kwan B, Dziedzic A, Jedlicka AE, Prigge ST. 2020. The NTP generating activity of pyruvate kinase II is critical for apicoplast maintenance in Plasmodium falciparum. eLife 9.

16. Maeda T, Saito T, Harb OS, Roos DS, Takeo S, Suzuki H, Tsuboi T, Takeuchi T, Asai T. 2009. Pyruvate kinase type-II isozyme in Plasmodium falciparum localizes to the apicoplast. Parasitol Int 58:101–105.

17. Zhang M, Wang C, Otto TD, Oberstaller J, Liao X, Adapa SR, Udenze K, Bronner IF, Casandra D, Mayho M, Brown J, Li S, Swanson J, Rayner JC, Jiang RHY, Adams JH. 2018. Uncovering the essential genes of the human malaria parasite Plasmodium falciparum by saturation mutagenesis. Science 360:eaap7847.

18. Zhong W, Li K, Cai Q, Guo J, Yuan M, Wong YH, Walkinshaw MD, Fothergill-Gilmore LA, Michels PAM, Dedon PC, Lescar J. 2020. Pyruvate kinase from Plasmodium falciparum: Structural and kinetic insights into the allosteric mechanism. Biochem Biophys Res Commun 532:370–376.

19. Dillenberger M, Rahlfs S, Becker K, Fritz-Wolf K. 2022. Prominent role of cysteine residues C49 and C343 in regulating Plasmodium falciparum pyruvate kinase activity. Struct Lond Engl 1993 30:1452–1461.e3.

20. Quansah N, Sarah C, Yamaryo-Botté Y, Botté CY. 2024. Complex Endosymbiosis II: The Nonphotosynthetic Plastid of Apicomplexa Parasites (The Apicoplast) and Its Integrated Metabolism. Methods Mol Biol Clifton NJ 2776:43–62.

21. Botté CY, Dubar F, McFadden GI, Maréchal E, Biot C. 2012. Plasmodium falciparum Apicoplast Drugs: Targets or Off-Targets? Chem Rev 112:1269–1283.

22. Tisnerat C, Dassonville-Klimpt A, Gosselet F, Sonnet P. 2022. Antimalarial Drug Discovery: From Quinine to the Most Recent Promising Clinical Drug Candidates. Curr Med Chem 29:3326–3365.

23. Fernandes JF, Lell B, Agnandji ST, Obiang RM, Bassat Q, Kremsner PG, Mordmüller B, Grobusch MP. 2015. Fosmidomycin as an antimalarial drug: a meta-analysis of clinical trials. 10.2217/FMB1560. systematic-review. Future Medicine Ltd London, UK. https://www.futuremedicine.com/doi/10.2217/FMB.15.60. Retrieved 22 April 2024.

24. Low LM, Stanisic DI, Good MF. 2018. Exploiting the apicoplast: apicoplast-targeting drugs and malaria vaccine development. Microbes Infect 20:477–483.

25. Kantele A, Mero S, Lääveri T. 2022. Doxycycline as an antimalarial: Impact on travellers’ diarrhoea and doxycycline resistance among various stool bacteria - Prospective study and literature review. Travel Med Infect Dis 49:102403.

26. Godinez-Macias KP, Chen D, Wallis JL, Siegel MG, Adam A, Bopp S, Carolino K, Coulson LB, Durst G, Thathy V, Esherick L, Farringer MA, Flannery EL, Forte B, Liu T, Godoy Magalhaes L, Gupta AK, Istvan ES, Jiang T, Kumpornsin K, Lobb K, McLean KJ, Moura IMR, Okombo J, Payne NC, Plater A, Rao SPS, Siqueira-Neto JL, Somsen BA, Summers RL, Zhang R, Gilson MK, Gamo F-J, Campo B, Baragaña B, Duffy J, Gilbert IH, Lukens AK, Dechering KJ, Niles JC, McNamara CW, Cheng X, Birkholtz L-M, Bronkhorst AW, Fidock DA, Wirth DF, Goldberg DE, Lee MCS, Winzeler EA. 2025. Revisiting the Plasmodium falciparum druggable genome using predicted structures and data mining. Npj Drug Discov 2:3.

27. Dantas TBV, Moura IMR, da Costa RP, de Souza GE, Severino RP, Consolaro HN, de Oliveira LF, Bonatto V, Cass QB, Guido RVC, de Sousa LRF. 2025. Tandem Mass Spectrometry and Bio-Guided Isolation of Secondary Metabolites With Antiplasmodial Activity From Dalbergia miscolobium Bark. Chem Biodivers e01449.

28. Wiedemar N, Hauser DA, Mäser P. 2020. 100 Years of Suramin. Antimicrob Agents Chemother 64:e01168–19.

29. Poigny S, Guyot M, Samadi M. 1998. One-step Synthesis of Tyromycin A and Analogues. J Org Chem 63:1342–1343.

30. Chepkirui C, Cheng T, Sum WC, Matasyoh JC, Decock C, Praditya DF, Wittstein K, Steinmann E, Stadler M. 2019. Skeletocutins A-L: Antibacterial Agents from the Kenyan Wood-Inhabiting Basidiomycete, Skeletocutis sp. J Agric Food Chem 67:8468–8475.

31. Amin PM, Su Z, Wang S. 2021. Gold-Catalyzed Cycloisomerization of Propargyl Pyruvates Enabling Unified Access to Tricladolides C and D, Chaetomellic Anhydride A, and Tyromycin A. J Org Chem 86:15318–15325.

32. Nagaraj VA, Sundaram B, Varadarajan NM, Subramani PA, Kalappa DM, Ghosh SK, Padmanaban G. 2013. Malaria parasite-synthesized heme is essential in the mosquito and liver stages and complements host heme in the blood stages of infection. PLoS Pathog 9:e1003522.

33. Vaughan AM, O’Neill MT, Tarun AS, Camargo N, Phuong TM, Aly ASI, Cowman AF, Kappe SHI. 2009. Type II fatty acid synthesis is essential only for malaria parasite late liver stage development. Cell Microbiol 11:506–520.

34. Shears MJ, MacRae JI, Mollard V, Goodman CD, Sturm A, Orchard LM, Llinás M, McConville MJ, Botté CY, McFadden GI. 2017. Characterization of the Plasmodium falciparum and P. berghei glycerol 3-phosphate acyltransferase involved in FASII fatty acid utilization in the malaria parasite apicoplast. Cell Microbiol 19:e12633.

35. Lindner SE, Sartain MJ, Hayes K, Harupa A, Moritz RL, Kappe SHI, Vaughan AM. 2014. Enzymes involved in plastid-targeted phosphatidic acid synthesis are essential for Plasmodium yoelii liver-stage development. Mol Microbiol 91:679–693.

36. Wiley JD, Merino EF, Krai PM, McLean KJ, Tripathi AK, Vega-Rodríguez J, Jacobs-Lorena M, Klemba M, Cassera MB. 2015. Isoprenoid precursor biosynthesis is the essential metabolic role of the apicoplast during gametocytogenesis in Plasmodium falciparum. Eukaryot Cell 14:128–139.

37. Yeh E, DeRisi JL. 2011. Chemical rescue of malaria parasites lacking an apicoplast defines organelle function in blood-stage Plasmodium falciparum. PLoS Biol 9:e1001138.

38. Pesaresi A. 2023. Mixed and non-competitive enzyme inhibition: underlying mechanisms and mechanistic irrelevance of the formal two-site model. J Enzyme Inhib Med Chem 38:2245168.

39. UniProt Consortium. 2023. UniProt: the Universal Protein Knowledgebase in 2023. Nucleic Acids Res 51:D523–D531.

40. Birkholtz L-M, Bastien O, Wells G, Grando D, Joubert F, Kasam V, Zimmermann M, Ortet P, Jacq N, Saïdani N, Roy S, Hofmann-Apitius M, Breton V, Louw AI, Maréchal E. 2006. Integration and mining of malaria molecular, functional and pharmacological data: how far are we from a chemogenomic knowledge space? Malar J 5:110.

41. Pizzi E, Frontali C. 2001. Low-complexity regions in Plasmodium falciparum proteins. Genome Res 11:218–229.

42. Saïdani N, Grando D, Valadié H, Bastien O, Maréchal E. 2009. Potential and limits of in silico target discovery - Case study of the search for new antimalarial chemotherapeutic targets. Infect Genet Evol J Mol Epidemiol Evol Genet Infect Dis 9:359–367.

43. Cheng T, Chepkirui C, Decock C, Matasyoh JC, Stadler M. 2019. Skeletocutins M–Q: biologically active compounds from the fruiting bodies of the basidiomycete Skeletocutis sp. collected in Africa. Beilstein J Org Chem 15:2782–2789.

44. Mangaleswaran S, Argade NP. 2001. A Facile Synthesis of Naturally Occurring Aminopeptidase Inhibitor Tyromycin A. J Org Chem 66:5259–5261.

45. Wubben TJ, Chaudhury S, Watch BT, Stuckey JA, Weh E, Fernando R, Goswami M, Pawar M, Rech JC, Besirli CG. 2023. Development of Novel Small-Molecule Activators of Pyruvate Kinase Muscle Isozyme 2, PKM2, to Reduce Photoreceptor Apoptosis. Pharm Basel Switz 16:705.

46. Guan M, Tong Y, Liu X, Dong D, Zhou Y. 2017. Enzyme Kinetic Assay to Measure the Activity of Tumor M2 Pyruvate Kinase in Breast Cancer Patients. Ann Clin Lab Sci 47:676–686.

47. Trager W, Jensen JB. 1976. Human malaria parasites in continuous culture. Science 193:673–675.

48. Ashraf K, Tajeri S, Arnold C-S, Amanzougaghene N, Franetich J-F, Vantaux A, Soulard V, Bordessoulles M, Cazals G, Bousema T, van Gemert G-J, Le Grand R, Dereuddre-Bosquet N, Barale J-C, Witkowski B, Snounou G, Duval R, Botté CY, Mazier D. 2022. Artemisinin-independent inhibitory activity of Artemisia sp. infusions against different Plasmodium stages including relapse-causing hypnozoites. Life Sci Alliance 5:e202101237.

